# DLPFC-tDCS unable to modulate mind-wandering propensity nor underlying functional or effective brain connectivity

**DOI:** 10.1101/2022.05.26.493632

**Authors:** Sean Coulborn, Davinia Fernández-Espejo

**Affiliations:** School of Psychology, University of Birmingham; Centre for Human Brain Health, University of Birmingham; University of California, Berkeley

## Abstract

There is conflicting evidence over the ability to modulate mind-wandering propensity with anodal transcranial direct current stimulation (tDCS) over the left dorsolateral prefrontal cortex (DLPFC-tDCS). Here, 20 participants received 20-minutes of anodal and sham DLPFC-tDCS while in the MRI scanner, in two separate sessions (counterbalanced). In each session, they completed two runs of a sustained attention to response task (before and during tDCS), which included probes recording subjective responses of mind-wandering. We assessed the effects of tDCS on behavioural responses as well as functional and effective dynamics, via dynamic functional network connectivity (dFNC) and dynamic causal modelling analyses over regions of the default mode, salience and executive control networks. Behavioural results provided substantial evidence in support of no effect of tDCS on task performance nor mind-wandering propensity. Similarly, we found no effect of tDCS on frequency (how often) or dwell time (time spent) of underlying brain states nor effective connectivity. Overall, our results suggest that DLPFC-tDCS is unable to modulate mind-wandering propensity or influence underlying brain function. This expands previous behavioural replication failures in suggesting that DLPFC-tDCS may not lead to even subtle (i.e., under a behavioural threshold) changes in brain activity during self-generated cognition.

## Introduction

Research investigating the use of transcranial direct current stimulation (tDCS) as a tool to modulate self-generated cognitive processes, such as mind-wandering, is becoming increasingly popular^1–11^. Mind-wandering is an internally directed process which is typically associated with the activation of the default mode network (DMN)^12,13^. This is a network found to be more active at rest and when engaging with decisions about oneself, with core regions including the posterior cingulate cortex / precuneus (PCC), the medial prefrontal cortex (MPFC), and the inferior parietal lobules^14^. Typically, DMN activity is thought to anticorrelate with that of so-called ‘task-positive’ networks such as the executive control network (ECN)^15^ or the salience network (SN). However, recent research has demonstrated that the dorsolateral prefrontal cortex (DLPFC), a key area in the ECN associated with high order cognitive functions^16^ and working memory^17^, also plays a key role in mind-wandering^18,19^. Based on this, an increasing number of studies target the DLFPC with non-invasive brain stimulation to try to modulate mind-wandering. In a series of studies, anodal left DLPFC-tDCS (with the return over the right supraorbital area) successfully increased self-reported mind-wandering propensity during a sustained attention to response task (SART) with interspersed thought probes^4,5^. This provided the first causative evidence for the role of DLPFC in mind-wandering. However, a recent pre-registered large-scale replication has called this conclusion into question by failing to elicit the same effects and instead reporting Bayesian support for the lack of an effect of tDCS stimulation^11^. Subsequent research reported further successful modulations of mind-wandering propensity after DLPFC-tDCS but in an inconsistent manner. For example, one study found cathodal (but not anodal) stimulation of the DLPFC increased mind-wandering compared to sham^10^, while a further study using a high-definition montage showed that anodal DLPFC-tDCS reduced mind-wandering propensity; both showing effects in the opposite direction to previously reported^20^. In contrast, and in line with the original findings, subsequent research reported increased mind-wandering propensity during anodal stimulation of the left DLPFC, but crucially, when the cathode was placed over the right inferior parietal lobule^6,7,9^. This leaves it unclear as to whether the DLPFC or the right inferior parietal lobule (and therefore the DMN) was driving these effects. Overall, the lack of congruous effects in the literature questions the role of the DLPFC in mind-wandering and the efficacy of tDCS in modulating it.

More broadly, tDCS is known for leading to inconsistent behavioural changes across participants and an array of cognitive functions^21,22^. It is however possible that, for some participants, tDCS elicits subtle changes in brain activation and/or connectivity that are not strong enough to drive changes in behaviour. Understanding effects at the neural level may thus help explain some of the inconsistent results found at the behavioural level. For example, previous fMRI studies using the SART^4,5,8–10^, arguably the most commonly used task in the literature measuring mind-wandering propensity and the potential for tDCS to modulate, confirmed that subjective episodes of mind-wandering elicit activation in the DMN, while occasions when the participant reports being on-task are related to activation of the ECN^18,23^. To our knowledge, there is no research investigating the effect of tDCS on brain activation during this task.

In addition to standard task-related activation, dynamic causal modelling (DCM), a technique that maps the causal influence brain regions have over each other (i.e., effective connectivity), may reveal more subtle underlying effects of stimulation on network connectivity. Indeed, previous research found modulations to the effective connectivity within and between the ECN and DMN following tDCS over the inferior parietal lobule during the SART^8^. However, to our knowledge, no research to date has explored this after DLPFC-tDCS.

Finally, dynamic functional network connectivity (dFNC), a method that allows one to observe reoccurring ‘brain states’ and measure their frequency and dwell time, has been successfully applied to identify key states during engagement with the SART^24^. Specifically, Denkova and colleagues^24^ observed five reoccurring ‘brain states’ over three key intrinsic brain networks (DMN, ECN and SN)^24^. While Denkova’s study did not include tDCS, it provides evidence that dFNC may be a sensitive method to detect changes in brain function associated with mind-wandering.

In this study, we used the SART with interspersed thought-probes before and during tDCS over the left DLPFC and concurrently during functional MRI acquisition to investigate whether tDCS could indeed modulate mind-wandering propensity as well as brain activation, and dynamic functional and effective connectivity. We expect to see increased activation of the DMN during episodes of mind-wandering and activation of the ECN when focussed on the SART task. If the left DLPFC is indeed causally involved in mind-wandering, we predict anodal stimulation will result in increased reports of mind-wandering and underlying activation of this region. In addition, we conducted exploratory analyses to investigate the effects of tDCS on effective connectivity and dFNC within and between the DMN, ECN, and SN.

## Method

### Pre-registration

We pre-registered this study at https://osf.io/8um5s/ on 16^th^ July 2019. Specifically, we pre-registered the design of the task, the stopping criteria for data collection, and the behavioural and GLM analysis. For the GLM analysis our report deviates away from the pre-registered pre-processing steps to instead use the increasingly more standard fMRIprep pipeline to allow for easier replicability. However, we performed the statistical analysis (GLM) as described in the pre-registration. Dynamic and effective connectivity analyses were exploratory and not pre-registered.

### Participants

27 right-handed healthy participants from an opportunity sample took part in the study, of which 26 completed both sessions (16 females, 10 males, aged 19-35, M = 24.50, SD = 3.56). We removed 6 participants who failed to respond to 10% or more of the SART trials or thought probes as specified in our pre-registration, as this indicates they were not following task instructions. This resulted in data from 20 participant’s being analysed (13 females, 7 males, aged 19-35, M = 24.85, SD = 3.92). We compensated participants with £30 for completing both parts of the study. The University of Birmingham’s Science Technology Engineering and Mathematics Ethical Review Committee provided ethical approval for the study. Before their inclusion in the study, we screened all participants for tDCS and MRI safety. All participants provided informed consent prior to their participation.

### Materials

#### Brain stimulation

We administered tDCS via an MRI compatible NeuroConn DC-Stimulator (neuroCare Group GmbH), using 5 × 5 cm rubber electrodes covered with Ten20® conductive gel. We placed the anode electrode over the left DLPFC and the cathode electrode over the right supraorbital area. The position of the electrodes was counterbalanced for the sham conditions. Before applying the electrodes, we cleaned the skin with alcohol wipes. To identify the location of let DLPFC on each participant and session, we used a 64-channel EEG cap, and marked F3 with a pen on the participant’s scalp. After that, we marked Fp2 to indicate the right supraorbital area. We removed the cap and fixed the electrodes in place (centred on the marks) using self-adhesive tape. Participants received 20 minutes of continuous 1.8mA stimulation with 10 seconds fade-in and fade-out periods. We monitored impedance and made sure it stayed below 15 kΩ. Figure 1 shows a simulation of the expected current generated through the brain in standard MNI space.

**Figure 1.**
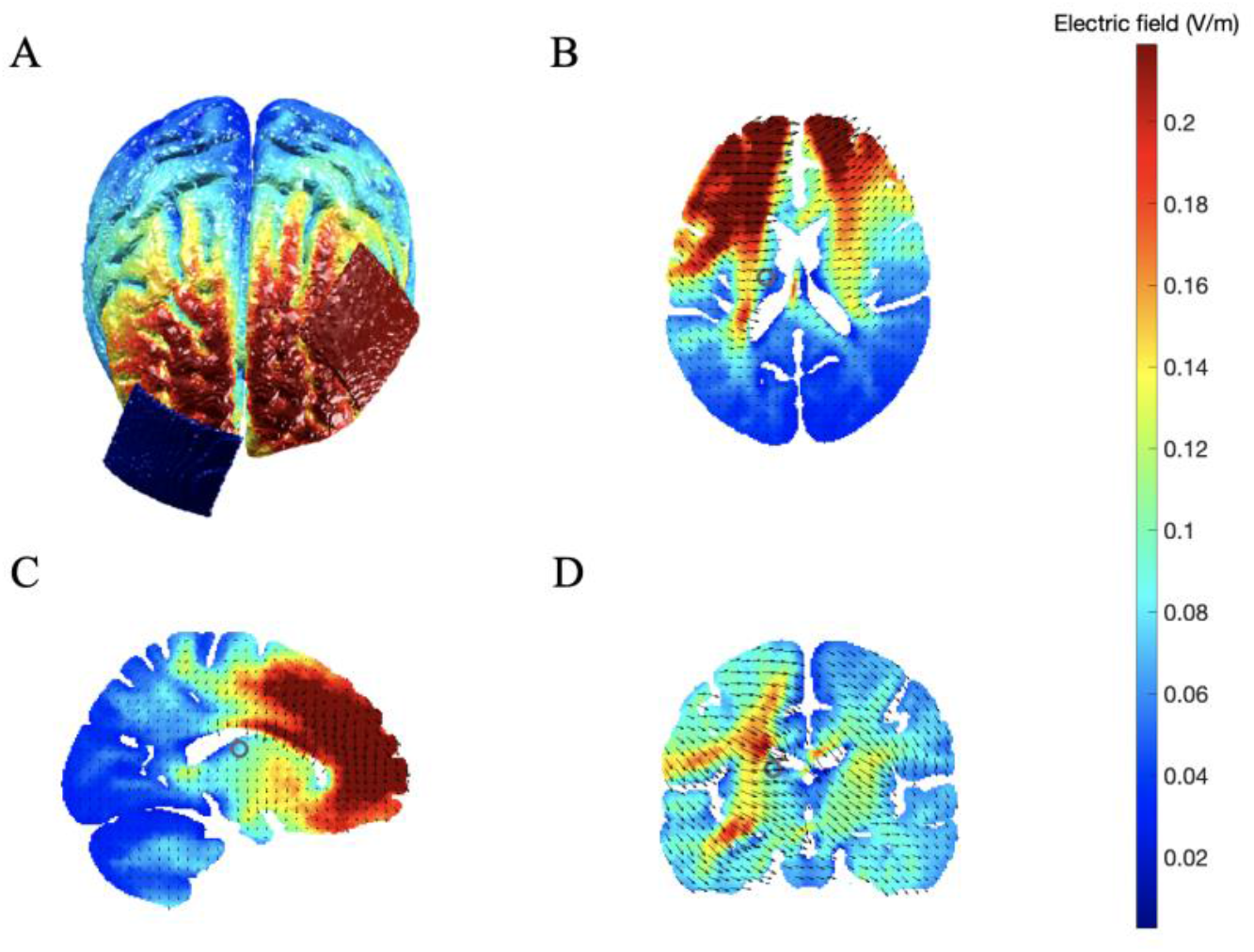
Computational modelof current magnitude and distribution produced with the open-source tool ROAST^25^. The simulation used the default ‘MNI152_T1_1mm’ template and was based on delivering 1.8mA current from the anode electrode (left DLPFC, F3), with the cathode placed over the right supraorbital area (Fp2). A) Illustration of the montage for left DLPFC (anode, red patch) and right supraorbital area (cathode, blue patch) and electric field distribution on the cortical surface. B) C) and D) show electric field strength and current flow in black arrows for axial, sagittal and coronal views, respectively. The colour chart on the right represents the magnitude of the electric field in Volts per meter. We used default conductivities used for the simulation (white matter 0.126 S/m; grey matter 0.276 S/m; cerebral spinal fluid 1.65 S/m; bone 0.01 S/m; skin 0.465 S/m; air 2.5e-14 S/m; gel 0.3 S/m; electrode 5.9e7 S/m).

#### Sustained Attention to Response Task (SART) and Thought-Probes

We used the SART, with similar parameters to Axelrod et al., 2015^5^, to elicit mind-wandering in a controlled manner (see Figure 2 for the experimental design). Specifically, participants responded with a button press to the visual presentation of a digit (from 0 to 9) as quickly and accurately as possible, except from the instances where the digit ‘3’ was presented (target) and the participant had to refrain from responding. Failure to do so would result in a commission error. Overall, the task included 535 non-targets (95.4% of trials) and 26 targets (4.6% of trials). Targets never appeared in the final 10s prior to a thought-probe. All digits were displayed for 1 second, followed by a fixation cross, also displayed for 1s. Throughout the task, we presented 32 thought-probes at pseudo-random intervals (20, 30, 40 and 50s; average 35s). These read ‘Where was your attention focused just before the probe?’. The participants responded by indicating whether they were ‘on-task’ or ‘off-task’. They had 4s to make a response before the SART began again. The task lasted for 20 minutes and 14 seconds.

**Figure 2.**
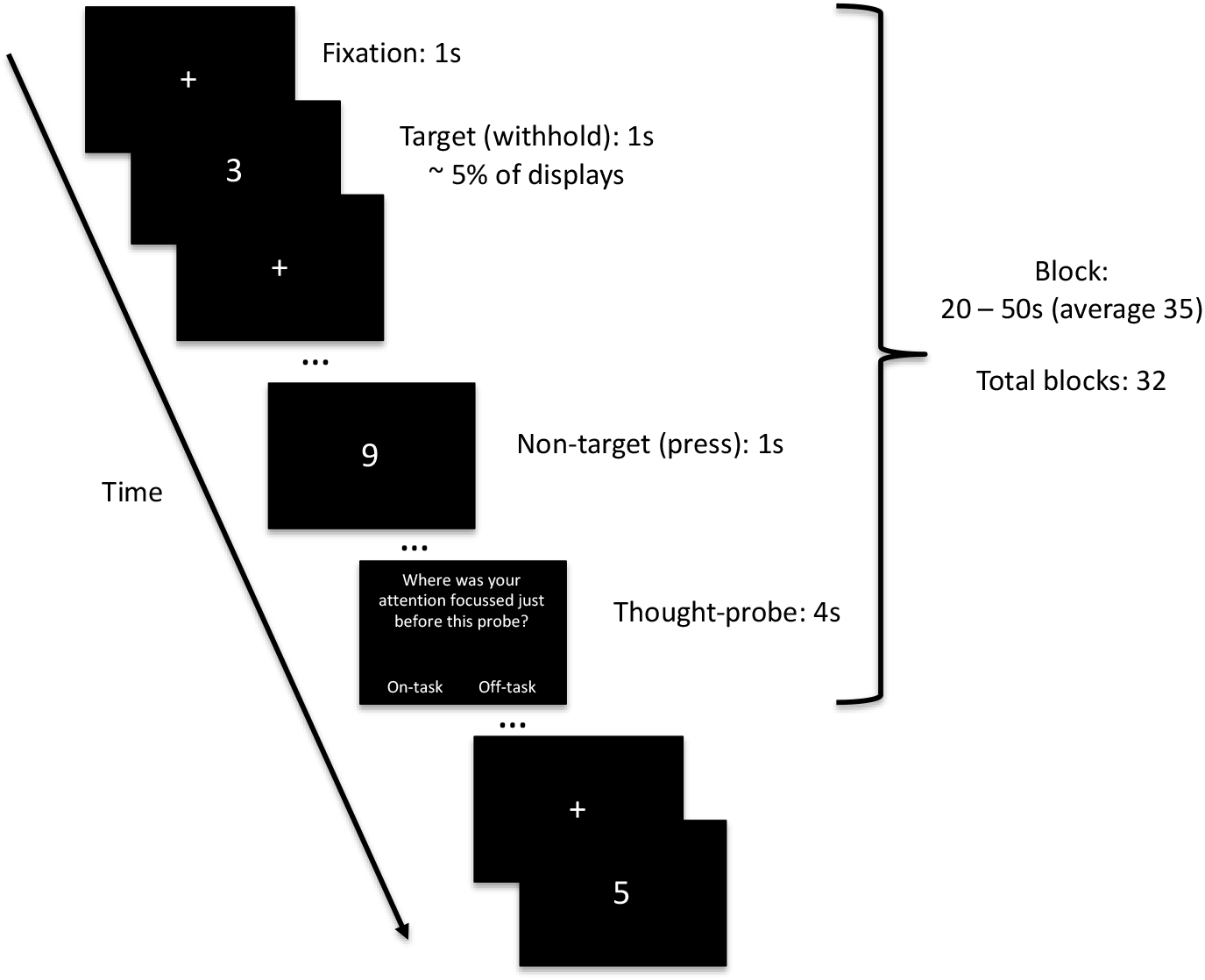
Flow diagram of the SART experimental design

### Experimental design & procedure

We adopted a within-subjects design with participants completing both conditions (anodal and sham) a minimum of 7 days apart (M = 10, SD = 3.87). Both sessions were identical except for the stimulation condition. The order of the anodal and sham sessions was counterbalanced, and participants were blind to the condition on each session.

Participants first completed three practice blocks (including SART and one thought probe per block) of the task outside of the scanner lasting approximately 30 seconds each to become familiar with the task. They then completed the Karolisnka sleepiness scale. Following this, participants entered the scanner and completed an offline (no tDCS) run of the SART, followed by an online (tDCS) run, which took place alongside 20 minutes of tDCS. The task was delivered using MATLAB 2017b and Psychtoolbox-3 on a Macbook Air (processor: Intel Core i7) for the practice run outside the MRI room, and MATLAB 2018a and Psychtoolbox-3 on a Windows 10 computer (processor: Intel Core i7) for the offline and online runs in the MRI scanner. At the end of each session, participants completed a post-tDCS questionnaire to assess their perceptions of the stimulation and again completed the Karolisnka sleepiness scale.

### fMRI acquisition

We collected data using a 3.0 Tesla Siemens PRISMA MRI scanner, using a 32-channel head-coil at the Centre for Human Brain Health (CHBH), University of Birmingham. We acquired a T1-weighted image (repetition time (TR) = 2 ms; echo time (TE) = 2.03 ms; matrix size = 256 × 256 mm; voxel size = 1 × 1 × 1 mm; field of view = 256 × 256 mm; and flip angle = 8 degrees) and echo-planar images with the following parameters: 57 slices (interleaved with multiband); TR = 1500 ms; TE = 35 ms; matrix size = 84 × 84 mm; voxel size = 2.5 × 2.5 × 2.5 mm; field of view = 210 × 210 mm; and flip angle = 71 degrees. This resulted in 812 volumes collected per run of the task.

We placed a NAtA Technologies Inc. response box (3 response buttons) into the participant’s right hand whilst in the scanner. The task and instructions were presented using a display back-projected onto a screen mounted at the end of the scanner. A mirror attached to the head coil enabled subjects to view the stimuli.

### Behavioural variables

We converted thought-probes of on-task and off-task responses into percentages per run of the task for analysis. We defined mind-wandering propensity as the percentage of off-task responses, as per previous research^7^. We also recorded reaction time after the presentation of non-target stimuli and calculated the mean reaction time per run of the task. We defined commission errors as a button press in response to a target and recorded the number of commissions errors per run of the task. We interpreted increased commission errors (to targets) and longer reaction times to non-targets as suggestions of greater levels of mind-wandering.

### fMRI pre-processing

We pre-processed the data using the standard pipeline on fMRIPrep (version 20.2.0)^26^, based on Nipype^27^. Specifically, anatomical T1 images were corrected for intensity and nonuniformity using N4BiasFieldCorrection^28^ distributed on ANTs 2.3.2^29^ and then skull-stripped them using antsBrainExtraction.sh workflow; we then segmented the brain tissue into cerebrospinal fluid (CSF), white-matter (WM) and grey matter (GM) using FAST on FSL v5.0.11, and finally spatially normalised to the ICBM 152 Nonlinear Asymmetrical template 2009c^30^. We then pre-processed functional images by first generating a reference volume (BOLD reference) and its skull-stripped version, again using the standard pipeline of fMRIPrep; then, we co-registered the BOLD reference to the T1w with boundary-based registration cost function^31^ configured with nine degrees of freedom accounting for the remaining distortion in BOLD images; following this, we estimated head motion parameters (transformation matrices and six corresponding rotation and translation parameters) before spatio-temporal filtering using *mcflirt*^32^ and performed slice timing correction, alongside motion-correction with additional ‘fieldmap-less’ distortion correction^33^; we then normalised the functional images into MNI space (MNI152NLin2009cAsym^30^); finally, we implemented framewise displacement^34^ (Power 2014) and DVARS^35^ on Nipype for each functional run.

We then performed spatial smoothing of the functional images in SPM12 (www.fil.ion.ucl.ac.uk/spm) on MATLAB version R2019b using an 8mm FWHM Gaussian kernel. To reduce low-frequency drift and high-frequency physiological noise for the dFNC and DCM analysis, we also applied a bandpass temporal filter between 0.01 - 0.1 Hz to the data using the DPABI toolbox in the DPARSF package (http://rfmri.org/DPARSF).

We removed one participant from all imaging analysis due to excessive head-movement across the scans (>15% of scans had > 0.9 framewise displacement in any given run of the task). We further removed a second participant from the univariate GLM analysis as they had no responses of ‘off-task’ in one of the runs of the task, and therefore this dataset would not allow for a valid comparison between ‘on-task’ and ‘off-task’ activity. The final sample was thus a total of 18 participants for the univariate analysis and a total of 19 for the dynamic and effective connectivity analyses.

### Brain activation

We estimated condition effects at each voxel according to a general linear model custom designed on SPM12 similar to that used by Christoff and colleagues^18^. Specifically, we used four regressors of interest: the first two comprised the 10s preceding each type of thought-probe response (‘on-task’ and ‘off-task’); we then created two further regressors including the 10s pre-target intervals preceding a commission error and those before a correct withhold, to represent occasions of being focussed on the task and being off task respectively. We also included regressors of no interest to model incorrect response to targets, correct response to targets and correct responses to non-targets, for the occurrence of each thought-probe and the response to a thought-probe, all with a duration of zero. This was to regress out activity associated with pressing a button to respond and to dissociate the occurrence of a thought-probe and the state of being on- or off-task. Finally, we included seven additional regressors capturing the six motion parameters estimated in the realignment step and the framewise displacement. Furthermore, we treated frames that exceed the threshold of >0.9mm framewise displacement as outliers and regressed them out using a stick regressor (as per Botvinik-Nezer et al., (2019)^36^).

For the first level analysis, we estimated pairwise contrasts between regressors of interest for each participant independently for offline and online runs. Specifically, we compared self-reported on-task and off-task responses. To obtain brain activity associated with performance in the task (as an indirect measure of mind-wandering) we compared commission errors and correct withholds to targets.

For the second level analyses, first we calculated a group level baseline activation using a 1-sample t-test for the offline run of the task taken during participants first session, in order to generate the canonical patterns of activation associated with our task and contrasts (on-task > off-task, off-task > on-task, commission errors > correct withholds and correct withhold > commission error). We also tested for the effects of stimulation using 2 (stimulation: anodal left-DLPFC, sham) x 2 (run: online, offline) repeated measure ANOVAs on each contrast above.

### Selection of regions of interest

We used a total of nine regions of interest (ROIs) for the functional connectivity analysis: two for the DMN (MPFC, PCC), four for ECN (bilateral DLPFC, bilateral PPC), and three for the SN (bilateral FIC and the ACC). To define these, we used the MNI coordinates used by Denkova and colleagues^24^ to center a sphere of 6mm radius (see Table 1). We then used these ROIs for the dynamic functional network connectivity and dynamic causal modelling analyses.

**Table 1.**
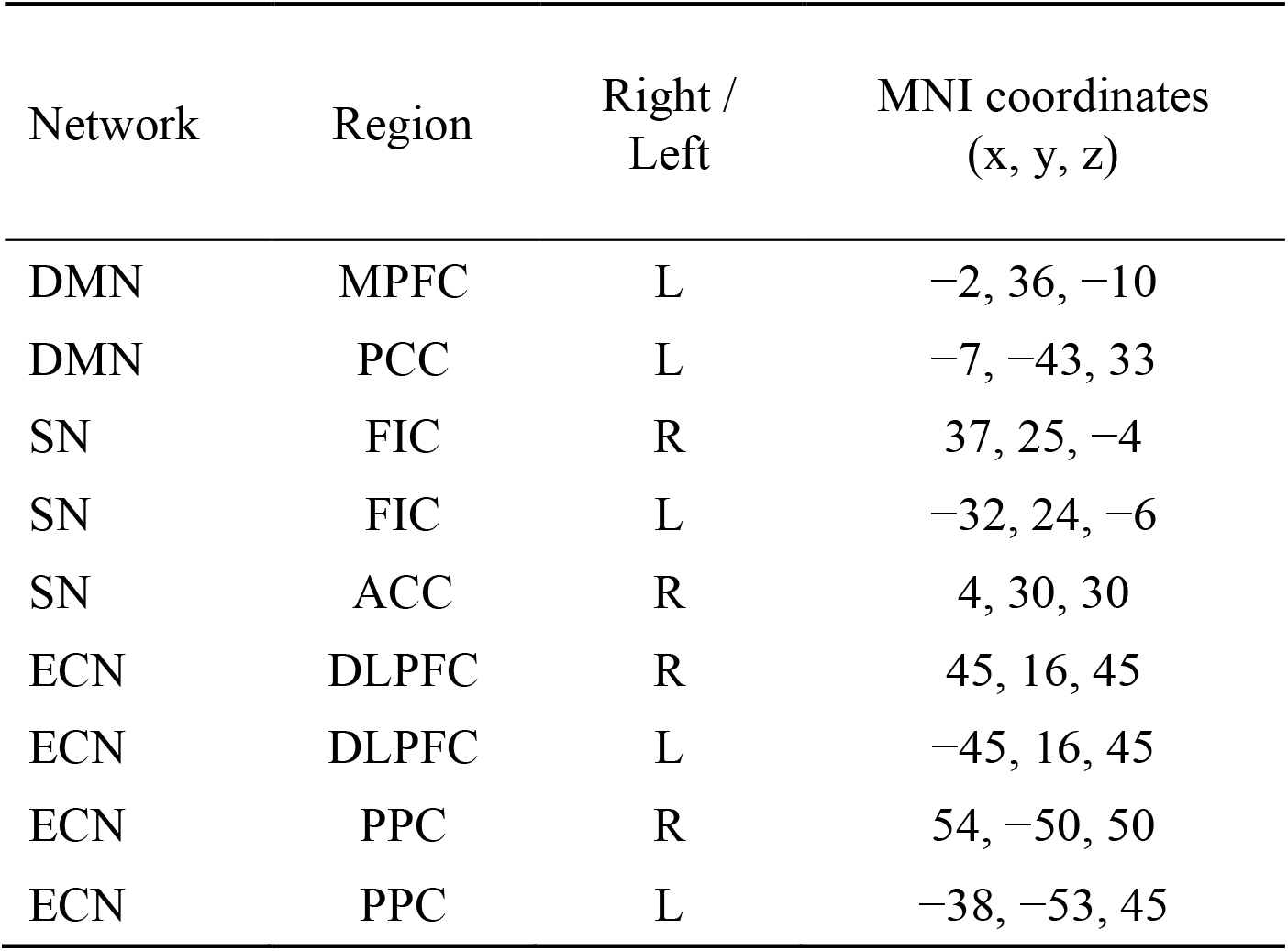
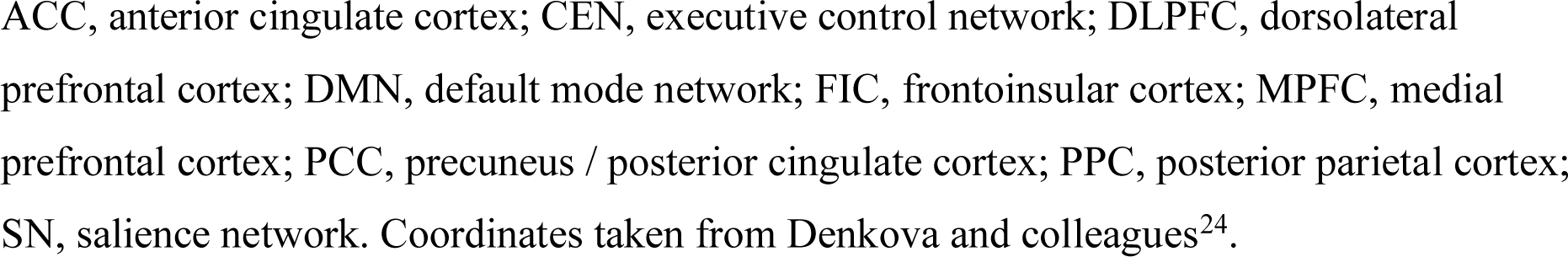
ROIs and MNI coordinates used in dFNC and DCM analyses. All had a sphere radius of 6mm.

| Network | Region | Right /<br>Left | MNI coordinates<br>(x, y, z) |
| --- | --- | --- | --- |
| DMN | MPFC | L | -2, 36, -10 |
| DMN | PCC | L | -7, -43, 33 |
| SN | FIC | R | 37, 25, -4 |
| SN | FIC | L | -32, 24, -6 |
| SN | ACC | R | 4, 30, 30 |
| ECN | DLPFC | R | 45, 16, 45 |
| ECN | DLPFC | L | -45, 16, 45 |
| ECN | PPC | R | 54, -50, 50 |
| ECN | PPC | L | -38, -53, 45 |

ACC, anterior cingulate cortex; CEN, executive control network; DLPFC, dorsolateral prefrontal cortex; DMN, default mode network; FIC, frontoinsula cortex; MPFC, medial prefrontal cortex; PCC, precuneus / posterior cingulate cortex; PPC, posterior parietal cortex; SN, salience network. Coordinates taken from Denkova and colleagues<sup>24</sup>.

### Dynamic functional network connectivity

After the functional images were smoothed, we regressed the six rigid body motion parameters and framewise displacement calculated during pre-processing. As above, we treated frames that exceed the threshold of >0.9mm framewise displacement as outliers and regressed them out using a stick regressor (as per ^36^). We followed the same pipeline and parameters used by Denkova and colleagues^24^. Specifically, we extracted time series for the nine ROIs and then submitted these for DFC analysis with a sliding window approach using the GIFT toolbox (https://trendscenter.org/software/gift/). We employed a window length of 44s and step size of 1 TR (1.5s) and did not isolate or delete probe periods as these are an integral part of the task. Therefore, the sliding windows focussed on an uninterrupted continuous temporal dynamic. This produced a correlation matrix which was 768 (sliding windows) x 36 (paired connections) per run per subject. We set the number of clusters (k) to 5 (as per^24^). We then calculated dFNC metrics for each run of the task per subject. These consisted of frequency of occurrence (percentage a brain state occurred) and dwell time (average time in sliding windows) spent in each of those 5 states. We also recorded the number of transitions (changes from one state to another). We describe the statistical analysis of these extracted measures below, alongside the behavioural data.

### Effective connectivity analysis

#### Spectral dynamic causal modelling

To assess effective connectivity, we employed a similar pipeline to that used by Kajimura and colleagues^8^. We initially ran a separate linear model to extract the time series to be used during dynamic causal modelling. This included the 6 rigid body parameters, framewise displacement as well as the cerebral spinal fluid and white matter signals as nuisance regressors. As above, we removed volumes with motion > 0.9mm framewise displacement using a stick regressor.

We then performed bilinear, one-state spectral DCM with the DCM12 routine on SPM12 using MATLAB version R2019b (see ^37^ for more details). We extracted time series from the same 9 ROIs as used in the dynamic functional connectivity analysis (see Table 1). We used a fully and reciprocally connected model resulting in a total of 81 effective connections (the A-matrix) produced and estimated for each model.

#### Parameter estimation and effects of tDCS on effective connectivity

Following model specification and estimation, we took the first level individual DCMs to the second level (between-subjects) using parametric empirical Bayes (PEB)^38^. PEB uses a hierarchical Bayesian framework to model individual subject’s effective connectivity relative to the group level by using individual DCMs (first level) as priors to constrain the variables in the Bayesian linear regression model (second level)^39^. Following this, we used Bayesian model reduction (BMR) to prune away any parameters not contributing to the evidence from the full model until no more improvements could be made. We then applied Bayesian model averaging (BMA)^38,39^ to take the remaining parameters, weigh them by their model evidence, and combine them into the final group model. We defined statistical significance by applying a threshold of posterior probability > 0.95 (strong evidence) for free energy.

To capture the typical group mean connectivity elicited whilst undertaking the task, we performed a group level PEB for participants’ first offline run. To test the effects of tDCS on the model parameters, we first created a PEB model in each participant which captured the pairwise interaction for greater increases during anodal stimulation as compared to sham (offline run < online run x anodal > sham condition). We then entered each subject’s PEB into a subsequent PEB that encoded the commonalities at the group level (mean).

### Statistical analyses of behavioural, dynamic functional connectivity

We used frequentist and Bayesian equivalent comparisons (with default priors) on JASP^40^ to test the effect of tDCS on behavioural data (i.e., ‘on-task’ responses and performance on SART) and dynamic functional states (i.e., frequency of occurrence, dwell time, and number of transitions). Specifically, we conducted repeated measures ANOVAs with stimulation (anodal, sham) and run (offline, online) as factors for each variable.

Similarly, we used a repeated measures ANOVA with stimulation (anodal and sham) and time (before and after tDCS) as factors, to test any possible effects of tDCS on the score from the Karolinska sleepiness scale.

Finally, to assess any potential differences in the perception of the sensations generated by stimulation between anodal and sham, we performed a Chi-Squared test for the association between received and perceived (real or sham) stimulation. We also analysed differences in perception of sensations caused by tDCS (intensity, discomfort, tingling, pain, burning, itching) using paired samples t-tests.

For the frequentist tests, we set the level of significance at p < 0.05. For the Bayesian test, we evaluated both the presence and the absence of an effect by comparing how different models explain the data given the factors of interest^41^. We used a Jeffrey-Zellner-Siow Bayes factor (JZS-BF10) to contrast the strength of the evidence for models reflecting the null and main effects and interactions^42^. For ease of interpretation, we include BF10 for those >1 and invert them as BF01 for those <1. In addition, for all ANOVAS, we also calculated a Bayes factor for the inclusion or exclusion of the variable of interest (BFinc, BFexcl) by comparing all models that exclude the interaction with all models that include it. A JSZ-BF between 0.33 and 3 is considered to be weak/anecdotal evidence for an effect; 3–10: substantial evidence; 10–100: strong evidence; >100: very strong evidence^43^.

## Results

### Behavioural

#### Mind-wandering responses and SART performance

While the data for percentage of ‘on-task’ responses, number of commission errors, and mean reaction time to non-targets violated tests for normality, we still employed ANOVAs as this is known to be robust enough even when all assumptions are not met^44^.

Frequentist analyses revealed no interaction between stimulation (anodal and sham) and run (offline / online) for any of the measures studied. Bayesian analyses provided further support for the lack of an effect of tDCS: substantial to strong evidence for the null as compared to the full model for ‘on-task’ responses, mean reaction time, and commission errors; strong evidence in support of excluding the interaction for ‘on-task’ responses and mean reaction time; and anecdotal evidence in support of excluding the interaction for commission errors. See Table 2 for descriptive statistics and Table 3 for statistical analyses.

**Table 2.**
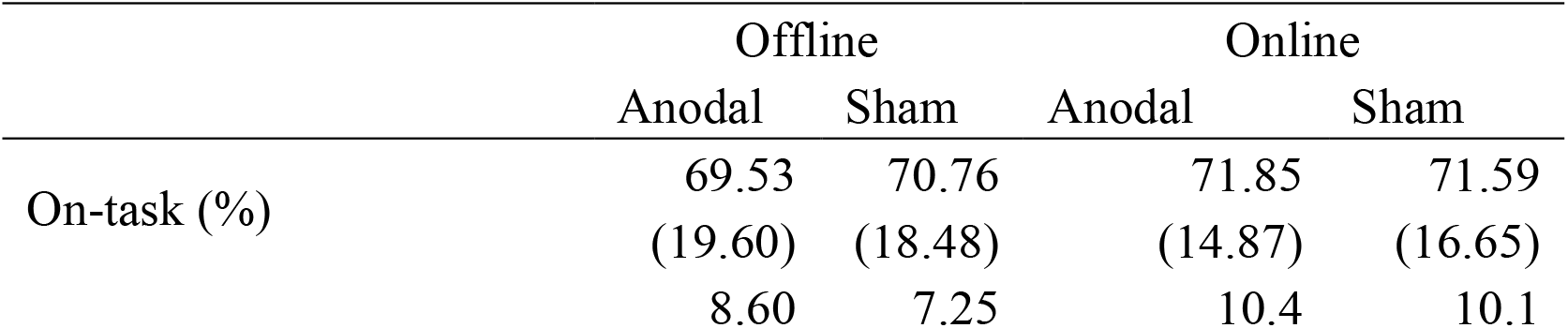
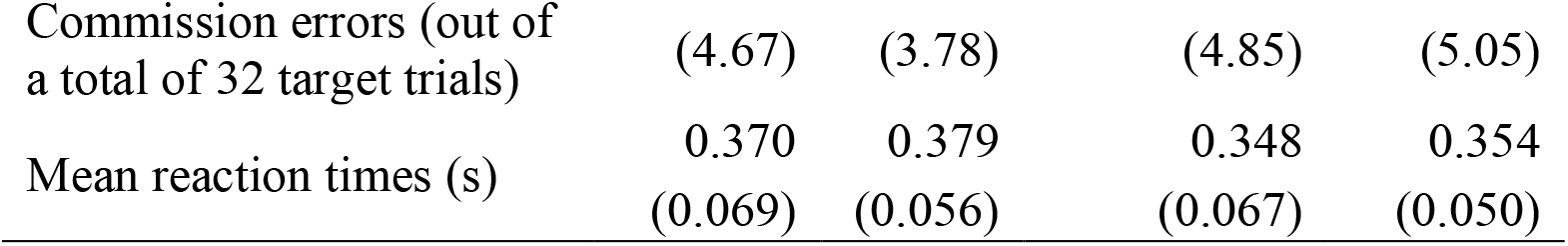
Percentage of ‘on-task’ responses to thought-probes and performance on SART. The data are group means (standard deviations in parenthesis).

**Table 3.** Bayesian and frequentists repeated measures ANOVA for the interaction of stimulation (anodal and sham) and run (offline and online) effects on subjective and objective measures of mind-wandering.

| | JZS-BF<sub>01</sub><br>for the null<br>vs full | BF<sub>excl</sub><br>(interact) | F | df | $\eta_p^2$ | P-value |
| --- | --- | --- | --- | --- | --- | --- |
| On-task (%) | 49.21** | 3.56* | 0.152 | 1,19 | 0.008 | 0.701 |
| Commission errors (%) | 4.42* | 2.31 | 0.929 | 1,19 | 0.929 | 0.347 |
| Mean reaction time (s) | 6.33* | 3.28* | 0.093 | 1,19 | 0.005 | 0.763 |

\*=BF > 3; \*\*=BF > 10

Similarly, we found no main effects for ‘on-task’ responses. However, we found a main effect of run for commission errors, F(1,19) = 22.06, p < 0.001, 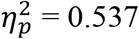; BF10 = 1 (BF10inc(run) = 108.124), with commission errors significantly higher in the online (second) run of the task. We also found a main effect of run for reaction times to non-targets, F(1,19) = 66.92, p < 0.001, 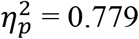; BF10 = 1 (BF10inc(run) = 399.692), with reaction times significantly faster during online runs.

As it can be appreciated in Figure 3, there was large variability across participants and sessions for both probe-responses and performance on the task.

**Figure 3.**
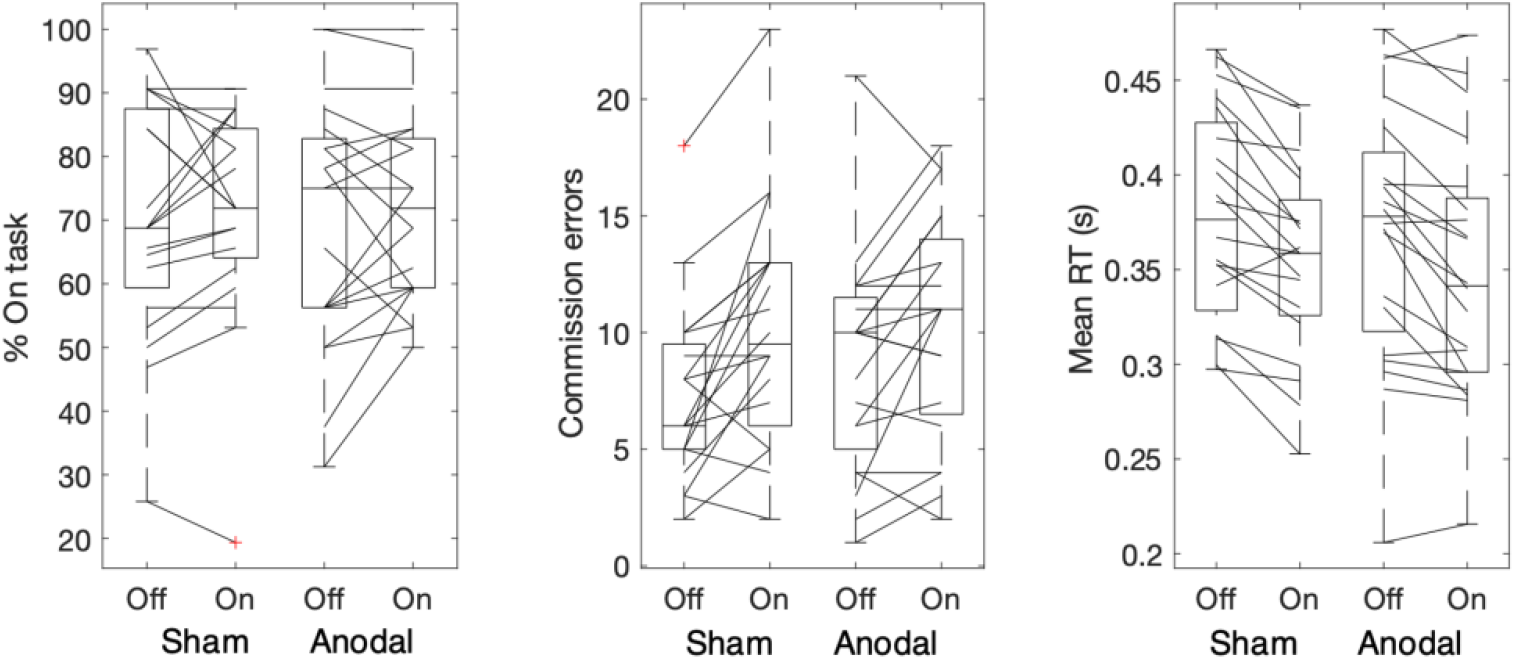
Line graphs with box plots displaying the variability in subjective responses (on-task) and variability in task performance (commission error and mean reaction time to non-targets) before (off) and during (on) tDCS in sham (left) and anodal (right) conditions. Outliers are indicated with a red cross. All measures exhibit large variability both at baseline and in response to tDCS.

#### Post-tDCS questionnaire

Table 4 displays the proportion of sessions where participants perceived to have had active stimulation from the post-tDCS perception questionnaire. A Chi-squared test revealed no significant association between stimulation type delivered and perception of receiving real or sham stimulation (X^2^(1) = 3.584 p = 0.058, BF10 (independent multinomial) = 2.379, N = 40).

**Table 4.**
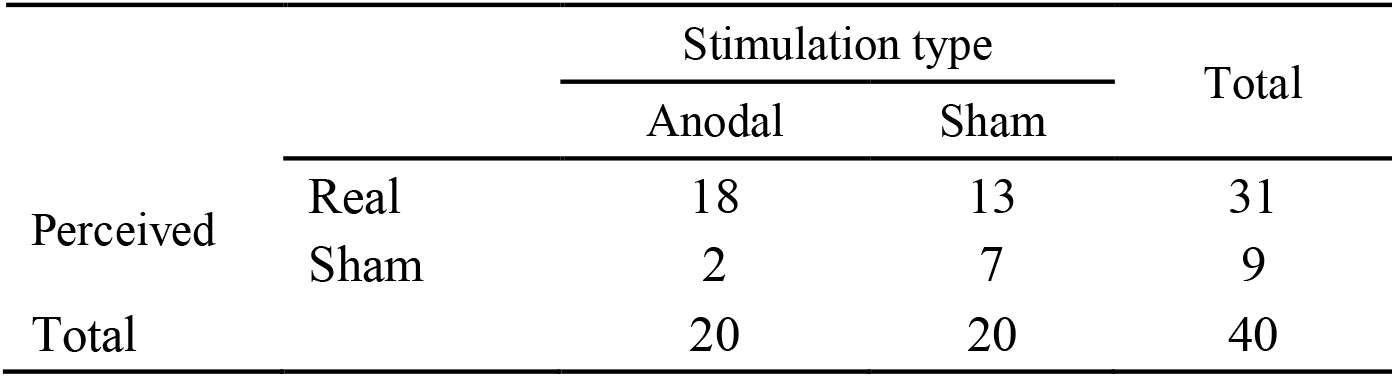
Contingency table displaying the frequency of perceived stimulation type (real or sham) against actual (anodal or sham).

Table 5 displays the results of the t-tests for additional perceptual scales. We found no significant differences for intensity, discomfort, tingling, pain, burning, and itching with substantial Bayesian support for lack of a difference for discomfort, pain, and itching (the rest were inconclusive). This suggests participants did not perceive a difference in sensations between stimulation conditions.

**Table 5.** t-tests for stimulation sensations from the post-tDCS questionnaire.

|  | t | df | p | BF<sub>10</sub> | BF<sub>01</sub> |
| --- | --- | --- | --- | --- | --- |
| Intensity | -1.894 | 19 | 0.074 | 1.031 | 0.97 |
| Discomfort | -0.358 | 19 | 0.724 | 0.246 | 4.061* |
| Tingling | -1.631 | 19 | 0.119 | 0.718 | 1.392 |
| Pain | -0.533 | 19 | 0.587 | 0.267 | 3.752* |
| Burning | -1.351 | 19 | 0.192 | 0.512 | 1.953 |
| Itching | -0.73 | 19 | 0.474 | 0.295 | 3.392* |

\*=BF > 3

#### Karolinska sleepiness scale

Both Bayesian and frequentist analyses revealed no interaction, with Bayesian analyses providing substantial support for the lack of an effect of tDCS for the null as compared to the full model as well as for all models excluding the interaction (F(1,19) = 0.053, p = 0.821, 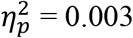; BF01 = 19.99, BF01excl(interaction) = 3.07). We observed a main effect of time (F(1,19) = 24.06, p < 0.001, 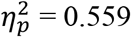; BF10 = 1 BF10inc(time) = 4431.12) with sleepiness scores significantly higher after the MRI sessions. See Figure 4 for summary statistics.

**Figure 4.**
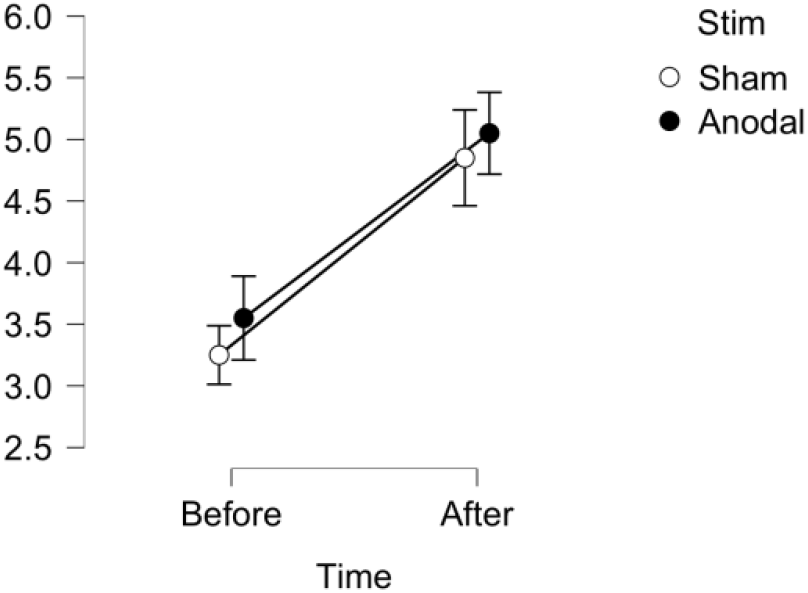
Mean scores for the Karolinska sleepiness scale (y-axis) before and after completion of the MRI session for both sham and anodal stimulation conditions. Bars represent standard error.

### Imaging

#### Baseline brain activity and connectivity

The univariate (GLM) analysis revealed no significant activations in the one-sample t-test capturing canonical activity on the offline run of the task corresponding to the first chronological session for any of our contrasts (‘off-task’ vs ‘on-task’, ‘on-task’ vs ‘off-task’, ‘commission error’ vs ‘correct withhold’, ‘correct withhold’ vs ‘commission error’) when corrected for multiple comparisons (p = 0.05 FWE).

**Table S1.**
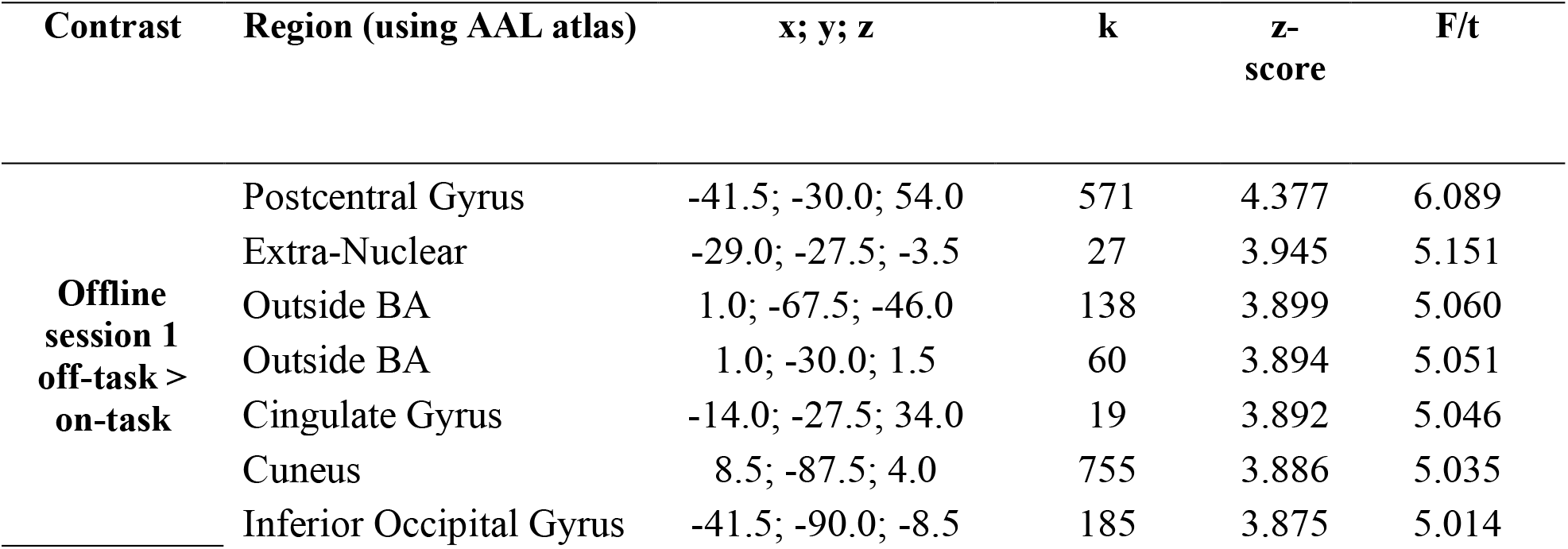
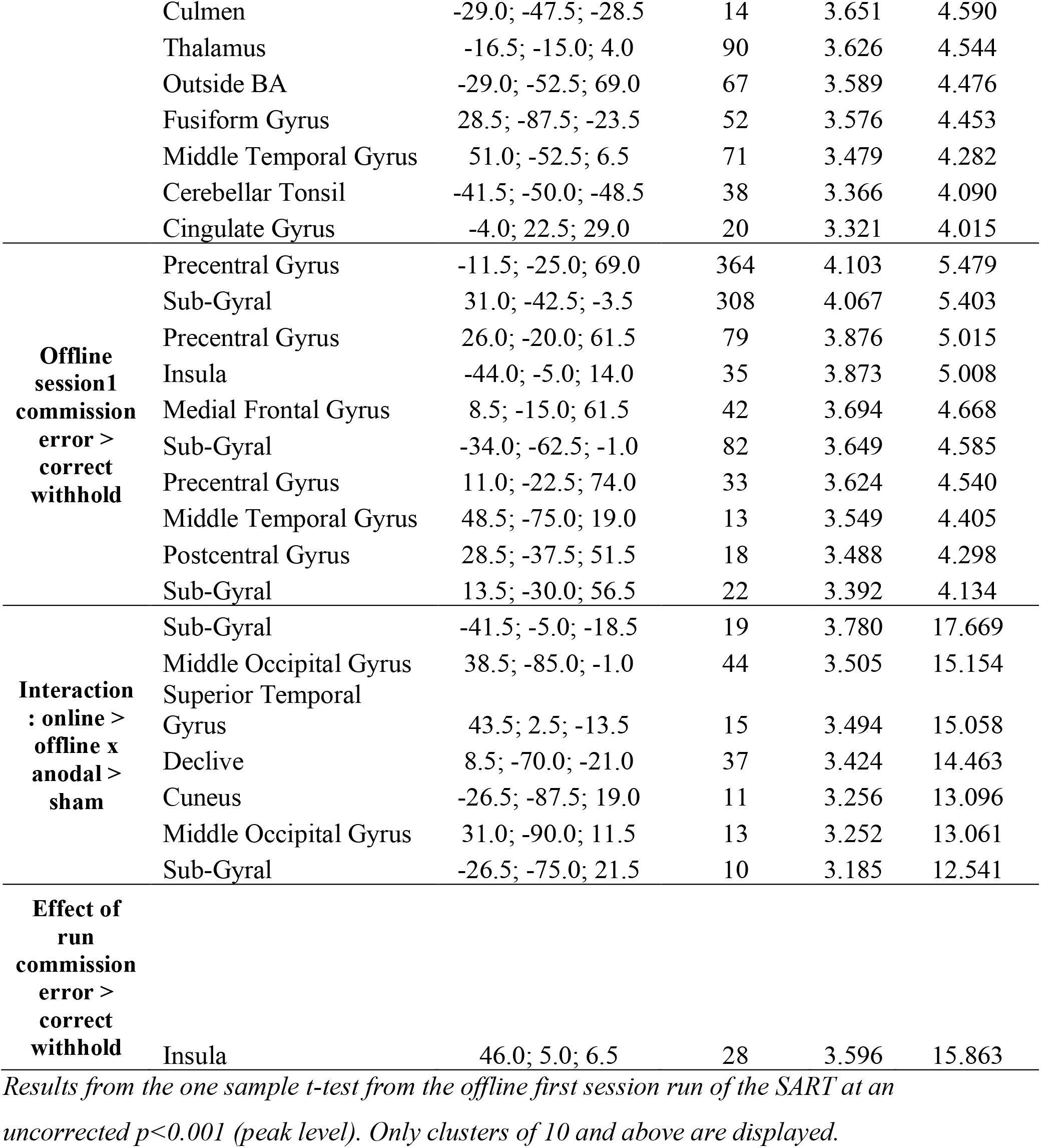
Baseline group activity.

| Contrast | Region (using AAL atlas) | x; y; z | k | z-score | F/t |
| --- | --- | --- | --- | --- | --- |
| **Offline session 1 off-task > on-task** | Postcentral Gyrus | -41.5; -30.0; 54.0 | 571 | 4.377 | 6.089 |
|  | Extra-Nuclear | -29.0; -27.5; -3.5 | 27 | 3.945 | 5.151 |
|  | Outside BA | 1.0; -67.5; -46.0 | 138 | 3.899 | 5.060 |
|  | Outside BA | 1.0; -30.0; 1.5 | 60 | 3.894 | 5.051 |
|  | Cingulate Gyrus | -14.0; -27.5; 34.0 | 19 | 3.892 | 5.046 |
|  | Cuneus | 8.5; -87.5; 4.0 | 755 | 3.886 | 5.035 |
|  | Inferior Occipital Gyrus | -41.5; -90.0; -8.5 | 185 | 3.875 | 5.014 |
|  | Culmen | -29.0; -47.5; -28.5 | 14 | 3.651 | 4.590 |
|  | Thalamus | -16.5; -15.0; 4.0 | 90 | 3.626 | 4.544 |
|  | Outside BA | -29.0; -52.5; 69.0 | 67 | 3.589 | 4.476 |
|  | Fusiform Gyrus | 28.5; -87.5; -23.5 | 52 | 3.576 | 4.453 |
|  | Middle Temporal Gyrus | 51.0; -52.5; 6.5 | 71 | 3.479 | 4.282 |
|  | Cerebellar Tonsil | -41.5; -50.0; -48.5 | 38 | 3.366 | 4.090 |
|  | Cingulate Gyrus | -4.0; 22.5; 29.0 | 20 | 3.321 | 4.015 |
| **Offline**<br>**session1**<br>**commission**<br>**error >**<br>**correct**<br>**withhold** | Precentral Gyrus | -11.5; -25.0; 69.0 | 364 | 4.103 | 5.479 |
|  | Sub-Gyral | 31.0; -42.5; -3.5 | 308 | 4.067 | 5.403 |
|  | Precentral Gyrus | 26.0; -20.0; 61.5 | 79 | 3.876 | 5.015 |
|  | Insula | -44.0; -5.0; 14.0 | 35 | 3.873 | 5.008 |
|  | Medial Frontal Gyrus | 8.5; -15.0; 61.5 | 42 | 3.694 | 4.668 |
|  | Sub-Gyral | -34.0; -62.5; -1.0 | 82 | 3.649 | 4.585 |
|  | Precentral Gyrus | 11.0; -22.5; 74.0 | 33 | 3.624 | 4.540 |
|  | Middle Temporal Gyrus | 48.5; -75.0; 19.0 | 13 | 3.549 | 4.405 |
|  | Postcentral Gyrus | 28.5; -37.5; 51.5 | 18 | 3.488 | 4.298 |
|  | Sub-Gyral | 13.5; -30.0; 56.5 | 22 | 3.392 | 4.134 |
| **Interaction**<br>**: online >**<br>**offline x**<br>**anodal >**<br>**sham** | Sub-Gyral | -41.5; -5.0; -18.5 | 19 | 3.780 | 17.669 |
|  | Middle Occipital Gyrus | 38.5; -85.0; -1.0 | 44 | 3.505 | 15.154 |
|  | Superior Temporal Gyrus | 43.5; 2.5; -13.5 | 15 | 3.494 | 15.058 |
|  | Declive | 8.5; -70.0; -21.0 | 37 | 3.424 | 14.463 |
|  | Cuneus | -26.5; -87.5; 19.0 | 11 | 3.256 | 13.096 |
|  | Middle Occipital Gyrus | 31.0; -90.0; 11.5 | 13 | 3.252 | 13.061 |
|  | Sub-Gyral | -26.5; -75.0; 21.5 | 10 | 3.185 | 12.541 |
| **Effect of**<br>**run**<br>**commission**<br>**error >**<br>**correct**<br>**withhold** |  |  |  |  |  |
|  | Insula | 46.0; 5.0; 6.5 | 28 | 3.596 | 15.863 |

*Results from the one sample t-test from the offline first session run of the SART at an uncorrected $p < 0.001$ (peak level). Only clusters of 10 and above are displayed.*

The dynamic functional connectivity analysis however, revealed 5 brain states that dynamically reoccurred during each run (i.e., when all sessions were pooled together regardless of stimulation and run). These are displayed in Figure 5. State 1 was characterised by slightly negative connectivity from DMN to both ECN and SN and a slight positive connectivity within networks. State 2 was characterised by slightly negative connectivity between DMN and SN, and DMN and ECN (specifically for the MPFC), with strong positive connectivity found between SN and ECN. State 3 and 4 were characterised by overall positive connectivity between and within networks, which was particularly stronger for state 4. State 5 was characterised by slight positive connectivity within and between networks, with the MPFC showing particularly weak connectivity with the rest of the regions.

**Figure 5.**
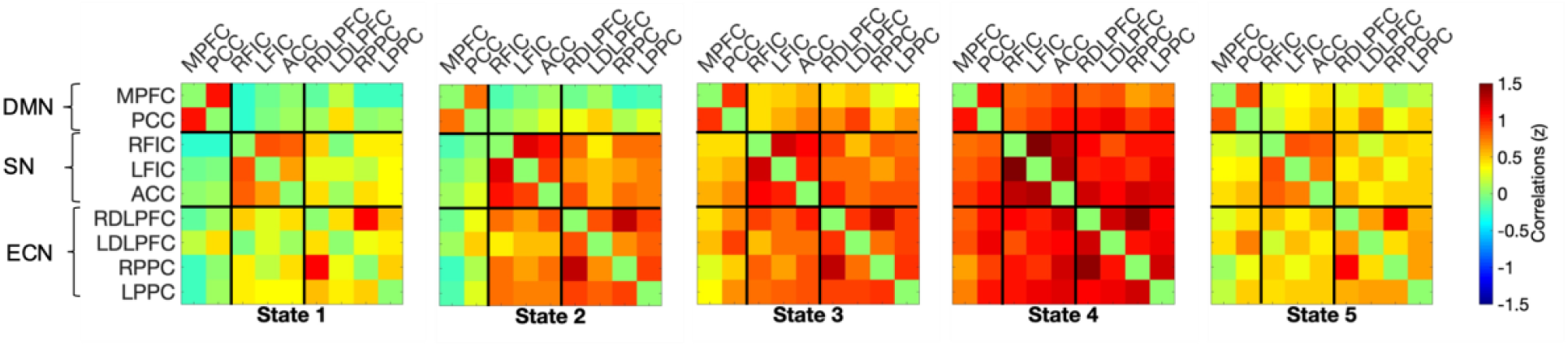
Five brain states which reoccurred during each session of the task. The colour bar on the right depicts the correlation strength (as z-score). ACC, anterior cingulate cortex; DLPFC, dorsolateral prefrontal cortex; DMN, default mode network; ECN, executive control network; FIC, frontoinsular cortex; MPFC, medial prefrontal cortex; PCC, precuneus / posterior cingulate cortex; PPC, posterior parietal cortex; SN, salience network.

Finally, Figure 6 shows the effective connectivity matrix depicting the group mean for the first session offline run of the task in the reduced model (only parameters with evidence above the threshold of 0.95 posterior probability). The DMN midline regions have an excitatory role over each other, and the PCC is strongly self-inhibitory. From the DMN to the SN, we observe excitatory coupling from the MPFC to both the right frontoinsular cortex and the anterior cingulate cortex, with inhibitory coupling from the PCC to the left frontoinsular cortex. From the DMN to ECN, the PCC exerted excitatory coupling towards bilateral DLPFC, while MPFC showed excitatory coupling with the left DLPFC and inhibitory towards right DLPFC and left posterior parietal cortex. The SN did not show any self-inhibitory connections but otherwise was largely excitatory within and between the networks. The ECN displayed excitatory coupling from both DLPFCs to PCC, and left DLPFC to MPFC, with both the posterior parietal cortices inhibitory towards MPFC. For the ECN to SN, we observed mixed communication with a combination of inhibitory and excitatory tones. For the ECN towards itself, we found strong self-inhibitory connections and an overall excitatory coupling within the network, except from left to right DLPFC which was inhibitory.

**Figure 6.**
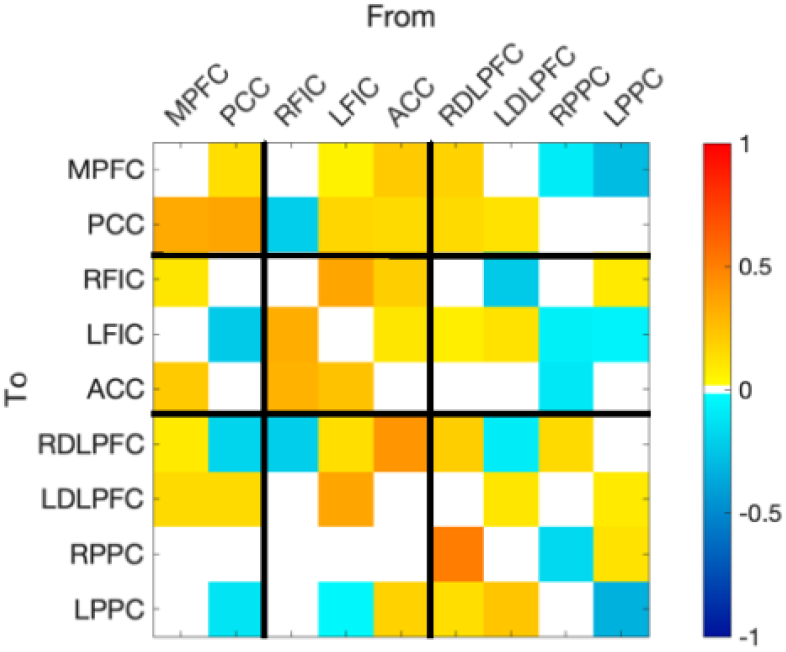
Group mean effective connectivity for session one offline run of the task. We only display connections present in the reduced model (>95% posterior probability for free energy). Extrinsic connections are represented in Hz and intrinsic self-connections are log scaled. The colour bar on the right indicates mean parameter strength (Ep.A). MPFC, medial prefrontal cortex; PCC, precuneus / posterior cingulate cortex; FIC, frontoinsular cortex; ACC, anterior cingulate cortex; DLPFC, dorsolateral prefrontal cortex; PPC, posterior parietal cortex; L, left; R, right.

#### Effects of tDCS on brain activity and connectivity

We did not observe any effects of tDCS (interaction), or a practice effect (main effect of run) on BOLD activation in the ANOVA. For exploratory purposes, we adopted a less conservative threshold of p < 0.001 (uncorrected), the results for which can be found in Supplementary Table 1.

In terms of brain states, both and Bayesian frequentist analyses revealed no interaction for any of the measures studied. Bayesian analyses indeed provided support for the lack of an interaction, suggesting no effect of tDCS. Specifically, we found strong evidence for the null as compared to the full model for frequency of states 1, 2 and 4, and dwell time in states 1-3, and found substantial evidence for all other measures (frequency of states 3 and 5, and dwell time for states 4-5). Furthermore, we found substantial evidence excluding the interaction for frequency of states 1, 2, 4 and 5, dwell time in states 1 and 2, and for the number of transitions, as well as anecdotal evidence for all other measures (frequency of state 3 and dwell time in states 3-5). Table 6 shows the results of the statistical analysis for the interactions. Figure 7 displays the descriptive statistics for frequency of occurrence and dwell time. We found a significant main effect of run for frequency of state 2 only, with increased occurrence in the online run compared to offline (see Table 7).

**Table 6.** Bayesian and frequentists repeated measures ANOVA for the interaction of stimulation (anodal and sham) and session (offline and online) on frequency, dwell time, and number of transitions.

| | JZS-BF<sub>01</sub><br>null vs full | BF<sub>excl</sub><br>(interact) | F | df | $\eta_p^2$ | P-value |
| --- | --- | --- | --- | --- | --- | --- |
| **Frequency** |  |  |  |  |  |  |
| State 1 | 30.601** | 3.118* | 0 | (1,18) | 0 | 0.992 |
| State 2 | 17.562** | 3.259* | 0.05 | (1,18) | 0.003 | 0.825 |
| State 3 | 5.889* | 1.545 | 1.685 | (1,18) | 0.086 | 0.211 |
| State 4 | 21.805** | 3.217* | 0.182 | (1,18) | 0.01 | 0.675 |
| State 5 | 9.468* | 3.060* | 0.286 | (1,18) | 0.016 | 0.6 |
| **Dwell Time** |  |  |  |  |  |  |
| State 1 | 13.641** | 3.304* | 0 | (1,18) | 0 | 0.979 |
| State 2 | 34.013** | 3.182* | 0 | (1,18) | 0 | 0.981 |
| State 3 | 22.821** | 1.891 | 0.963 | (1,18) | 0.051 | 0.34 |
| State 4 | 6.705* | 1.469 | 1.638 | (1,18) | 0.071 | 0.257 |
| State 5 | 9.508* | 2.435 | 0.632 | (1,18) | 0.034 | 0.437 |
| **Number of transitions** |  |  |  |  |  |  |
| All | 37.092** | 3.201* | 0.061 | (1,18) | 0.003 | 0.808 |

\*=BF > 3; \*\*=BF > 10

**Table 7.** Bayesian and frequentists repeated measures ANOVA for the main effect of run on frequency, dwell time, and number of transitions.

| | JZS-BF<sub>01</sub><br>null vs full | BF<sub>excl</sub> (run) | F | df | $\eta_p^2$ | P-value |
| --- | --- | --- | --- | --- | --- | --- |
| **Frequency** |  |  |  |  |  |  |
| State 1 | 2.238 | 1.267 | 1.153 | (1,18) | 0.060 | 0.297 |
| State 2 | 1.263 | 2.25 | 5.025 | (1,18) | 0.218 | 0.038<sup>#</sup> |
| State 3 | 4.644* | 2.25 | 0.343 | (1,18) | 0.019 | 0.566 |
| State 4 | 2.249 | 2.249 | 1.254 | (1,18) | 0.065 | 0.278 |
| State 5 | 1.000 | 0.505<br>(BF<sub>10</sub>=1.981<br>) | 3.690 | (1,18) | 0.016 | 0.600 |
| **Dwell Time** |  |  |  |  |  |  |
| State 1 | 7.851* | 1.405 | 0.089 | (1,18) | 0.005 | 0.769 |
| State 2 | 4.118* | 1.405 | 0.085 | (1,18) | 0.005 | 0.774 |
| State 3 | 2.921 | 1.405 | 0.818 | (1,18) | 0.043 | 0.387 |
| State 4 | 1.410 | 1.405 | 2.783 | (1,18) | 0.134 | 0.113 |
| State 5 | 1.000 | 0.876<br>(BF<sub>10</sub>=1.141<br>) | 0.632 | (1,18) | 0.034 | 0.437 |
| **Number of transitions** |  |  |  |  |  |  |
| All | 3.089* | 3.056* | 0.894 | (1,18) | 0.047 | 0.357 |

\*BF<sub>01</sub> > 3, <sup>#</sup>p < 0.050

**Figure 7.**
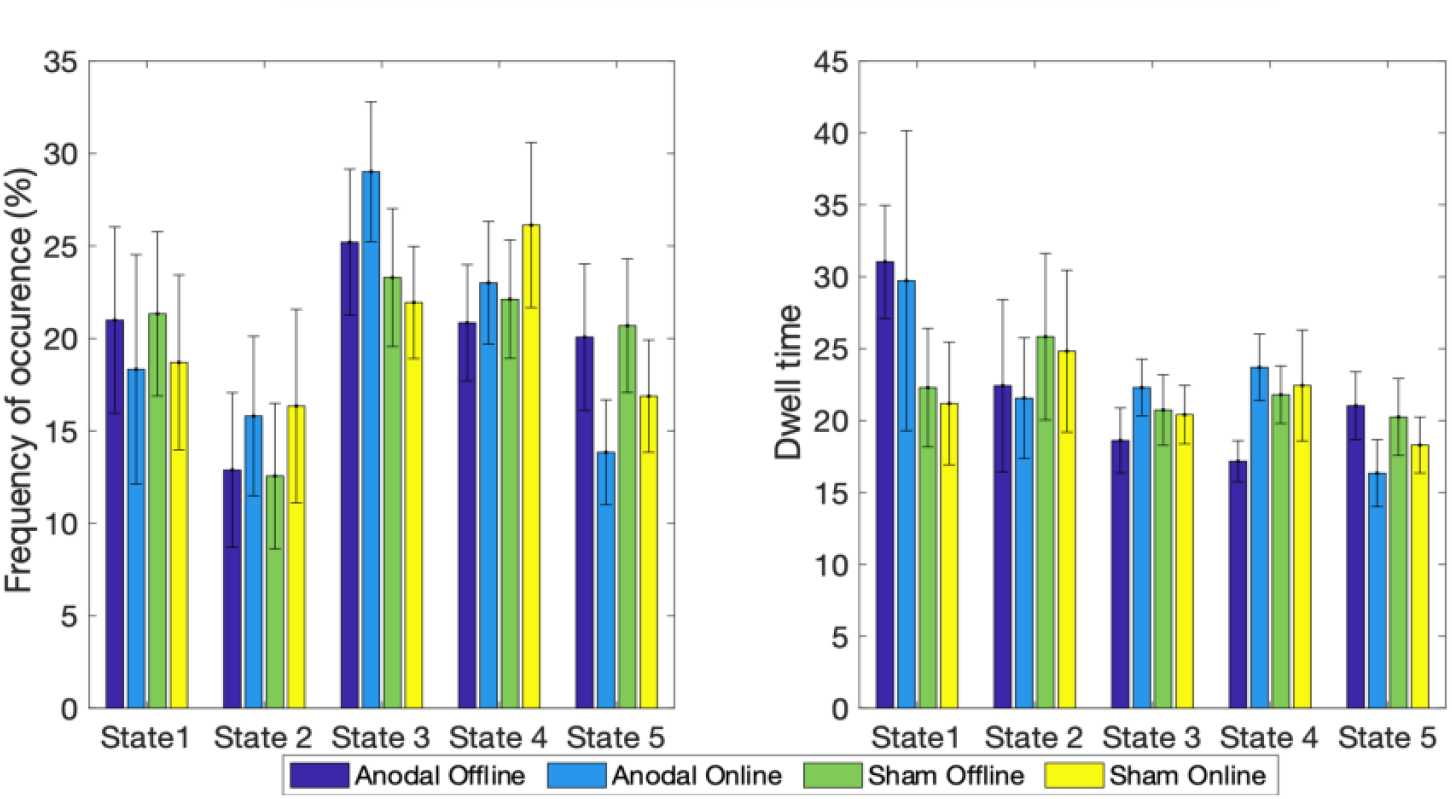
Frequency of occurrence and dwell time for each state and session. We found no significant interactions or main effects. Bayesian results support the lack of an effect of tDCS. Error bars represent standard error of the mean.

Similarly, the PEB modelling the interaction did not find evidence for an effect of tDCS across any of the parameters.

## Discussion

Here we investigated the efficacy of left DLPFC-tDCS in modulating mind-wandering propensity on a comparable sample size to previous research^4,6,7^ but using a within-subjects design to better control for individual differences. Furthermore, alongside behavioural metrics, we assessed the effects of tDCS on intrinsic brain activity as well as dynamic functional and effective connectivity. DLPFC-tDCS was unable to modulate subjective reports of mind-wandering, commission errors or reaction time in a SART task, underlying brain activation, frequency, dwell time or transitions across underlying functional states, nor effective connectivity across key underlying networks. Indeed, we provide Bayesian evidence supporting the lack of an effect of stimulation in most of these measures. Overall, our results therefore suggest, against our hypothesis, that left DLPFC-tDCS is unable to modulate mind-wandering propensity or underlying brain function.

Specifically, we showed strong Bayesian evidence for the lack of an effect of stimulation on subjective reports of on-task responses, and substantial evidence for the lack of an effect on commission errors and mean reaction time to SART, all when comparing the null versus full models. Furthermore, when considering the effect of the interaction itself, we found substantial evidence for the models excluding the interaction for on-task responses and mean reaction time (with anecdotal evidence for commission errors). Our behavioural results contradict early reports of successful modulations using this montage^4,5^ and instead support a recent multi-centre failed replication^11^. There are a few differences between our study and previous reports in the literature that should be acknowledged. First, we used an increased number of thought probes per run (32 compared to 12^5^ and 18^4^). However, recent research which varied the probe rate found no effect on probe responses caught by online sampling, and therefore our additional thought probes unlikely explain our reported lack of effect of tDCS^45^. Additionally, we used a higher intensity of stimulation (1.8mA compared to 1mA) to try to enhance our effects. While increasing current has sometimes been related to contradictory effects for cathodal stimulation^46^, it is typically acknowledged that this is not the case for anodal stimulation, where it should instead enhance the effects. Crucially, a recent study stimulating the DLPFC observed similar levels of increased mind-wandering after both 1mA and 2mA anodal stimulation of the DLPFC, as compared to sham^9^, further suggesting that our higher polarity is unlikely to have caused the lack of successful modulations in our study. It should be noted however that Filmer and colleagues placed the return electrode over the right inferior parietal lobule and therefore it is unclear which region is driving the effects in their case^9^. Future research is needed to understand the impact of current intensity and its potential role to help explain the observed discrepancies across the literature.

Recent reports have called into question the effectiveness of sham and suggested that placebo effect may have played a role in some successful tDCS reports^47,48^. Our results suggest participants were unable to discriminate between active or sham stimulation, nor showed any differences in their reported physical sensations. To further ensure any potential effects of external variables, we also measured sleepiness using the Karolinska Sleepiness Scale. Similarly, active vs sham tDCS did not have an effect on how tired the participants felt after completing the scan, suggesting that our lack of effects were not produced by differences in tiredness. We, however, found a strong effect of time, with sleepiness being significantly greater after completing the session. This was expected given the monotonous nature of the task and time spent lay down in the scanner and may explain the reduced reaction times and increased errors in the second run for each session. In addition to the lack of behavioural responses to tDCS, we were unable to observe any modulation to brain activation at the level of BOLD response, functional dynamic connectivity over time (in the form of frequency and dwell time of brain states, as well as number of transitions across states) or to effective connectivity within and between the DMN, EC and SN (any parameters within or between networks). In the case of dynamic connectivity, we indeed found evidence for a lack of an effect across most measures. To date, we are unaware of any published research investigating the effects of DLPFC-tDCS on brain activation specifically related to mind-wandering. There are a small number of studies investigating the effects of DLPFC-tDCS on brain activation in relation to executive function^49,50^. Two of these observed modulations in activation and functional connectivity of the DLPFC, albeit interestingly without any reported effects of tDCS on behaviour^49,50^. An important difference between our study and these is the tasks participants completed while in the scanner, which in their case were the Balloon Analogue Risk Task (a reward versus loss, risk-behaviour task) and the N-back task, both of which have been found to consistently recruit activation of the DLPFC. It is possible that, as these tasks repeatedly engage the target area (N-back^51^ and Balloon Analogue Risk Task^52^), observing any subtle changes during and post stimulation may be more likely. Indeed, it is becoming increasingly accepted that the effects of tDCS are modulated by the participants cognitive or brain state during stimulation^53^. Our participants were completing the SART while receiving tDCS, and thus it is expected that their thoughts would have been drifting between internally (mind-wandering) and externally (the actual SART) directed attention. This can indeed be observed in their subjective reports to thought probes. These drifts would have been accompanied by changes in the underlying brain state. As can be appreciated in Figure 3, we observed a large inter-subject variability for baseline levels of being on-task. This individual preference for being more on or off-task may thus have affected the extent to which each participant was in the most optimal state to be influenced by tDCS and explain the discrepancy between our results and previous research. In fact, individual differences would particularly affect those studies which used a between subjects’ design, as some cohorts may have contained more participants likely to mind-wander and perhaps more likely to benefit from the effects of stimulation.

Early fMRI research (without tDCS) demonstrated that the SART used with interspersed thought-probes successfully activated the DMN during reported episodes of ‘off-task’ and the ECN for ‘on-task’ responses^18,23^. However, subsequent research using the same design was unable to replicate this^24^ or used independent component analysis (ICA) to identify networks of interests instead^54^. Our results are in line with this and may suggest a lack of power to reliably identify the regions of interest in the GLM analyses, but also may highlight the unreliability of participant self-reports. Specifically, the thought probes require participants to subjectively identify whether they were on or off-task, and this is known to be problematic for some^55^, given the sometimes vague and all-encompassing classification of ‘mind-wandering’ thoughts^56^. Despite this, the SART remains the most typically used task in studies investigating mind wandering^18,24,57,58^, and in particular in conjunction with tDCS^1,4,5,8–11^. It is worth noting that we (and others) also analysed objective measures that do not rely on participants’ self-report (i.e., commission errors and reaction times) and can be used as proxies for mind-wandering propensity, but these also failed to elicit the expected brain activation or to be modulated by our tDCS montage.

Mind-wandering is a self-generated, internally directed process with propensity and characteristics unique to each individual. An ability to modulate this multifaceted self-referential process with an external medium, such as tDCS, would be an invaluable tool to researchers and clinicians. Manipulating participants’ propensity to mind-wander with such a technique via electrode placement over specific brain areas would help unveil the underlying neural processes involved in this complex phenomenon and reveal more about the causal influences of tDCS on brain activation. Mind-wandering, particularly an excessive propensity to do so, has been associated with poor task performance^59^ and with a number of neurological disorders (including: schizophrenia^60^, and depression^61^). Due to this relationship, DLPFC-tDCS has been posed as a potential therapeutic tool in these patient groups^62,63^. Our results, however, and in particular the lack of effects on brain activity and connectivity, call into question its ability to modulate this self-generated process and its use in clinical settings. An overarching concern surrounding tDCS research, and another possible explanation for our lack of effects, is that the currents typically used (<2mA) do not reach the brain with enough intensity to elicit any effects. Research in mice has suggested that a current of 1 V/m gradient is required to garner online effects on neuronal spiking^64^. In the current study we used 1.8mA stimulation on the surface, which translated into a predicted maximum cortical intensity of 0.235 V/m (based on the ROAST simulation, see Figure 1). It is thus possible that the delivered current was insufficient to perturb neuronal firing, hence explaining the lack of an effect on functional and effective connectivity. Despite this, previous research reported successful behavioural modulations using much lower amplitudes (e.g., 1mA^4,5^). Potential inter-individual variability in brain structure and the resulting differences in current flow across the brain for a population-level dose level^65^ may also help explain the discrepancies in reported results. Future research may benefit from implementing higher currents (e.g., ∼4mA, which has been shown to be safe in human subjects^66^), or more specifically tailored current dosage dependent upon individual anatomy and tissue properties, in order to reach the required current at the cortical level to elicit the desired neuronal changes.

Leaving the effects of tDCS aside, we partially replicated the brain states previously observed between DMN, ECN, and SN networks in research comparing connectivity during resting state fMRI and throughout the SART^24^. Specifically, States 3 and 4 were characterised by overall strong positive connectivity within and between networks and were the most frequent states throughout the task. We also we identified a state (State 1) showing the weakest overall connectivity within and between each network and may be associated with transitioning between states of focus and unfocussed attention^67,68^. State 2 showed negative connectivity between DMN and SN / ECN (more prominent for MPFC) and positive connectivity between SN and ECN. The SN has previously been associated with switches between the DMN and ECN to inhibit task-irrelevant self-directed thoughts and encourage task-relevant behaviour^69–71^. Denkova and colleagues^24^ found this state to be more frequent during task conditions compared to rest, and further research observed it during an attention-to-breath task suggesting it reflects a state of focussed attention^72^. Finally, State 5 displays a small but overall positive connectivity within and between networks (slightly less for the MPFC to SN and ECN) and appears to be the transitory phase between states 3 and states 1 and 2. Throughout all of the states we observe a stronger connectivity between the PCC and the DLPFC (specifically the left) compared with other regions outside the DMN. This supports research detailing the importance of this connection in mind-wandering and goal-directed cognition^73,74^ and could be reflecting the important role of the PCC being a major hub in switching between intrinsic networks^75–77^. Interestingly, we did not observe one of the states previously reported (state 2 in ^24^). This state was characterised by negative connectivity between SN and ECN, specifically for bilateral frontoinsular cortex, with connectivity being negative between the PCC and SN and positive between PCC and ECN. This discrepancy between our findings is not unexpected given that they report this brain state being most common during resting state (which we did not acquire in our study) as compared to throughout the SART, suggesting that this state is something more specifically characteristic at rest.

Regarding baseline levels of effective connectivity, we observed an overall excitatory coupling between regions within each network, in line with previous research^78^. Within the DMN, and in line with our dynamic functional connectivity results, we observe excitatory coupling in both directions between the PCC and MPFC. This supports previous research at rest^79^ and during cognitively demanding tasks (e.g., studies reporting a bi-directional inhibitory relationship during the N-back task^80^). Our results may indicate that the SART indeed elicits more self-referential thought processes resulting in these networks acting similar to when mind-wandering at rest. However, the literature here is inconsistent, with healthy controls previously showing an inhibitory role of the PCC towards the MPFC at rest^81,82^ and excitatory coupling during a gender judgement task^82^. At a cross-network level, from the DMN we see an inhibitory role of the PCC and an excitatory role of the MPFC towards both the SN and ECN. The excitatory coupling between the MPFC and the SN has been noted before, particularly towards the anterior cingulate cortex^78^. The DMN and ECN / SN have previously been reported to be anti-correlated^15^. Therefore, the excitatory coupling from the PCC to the MPFC in conjunction with the inhibitory coupling from the PCC to both ECN and SN may signify the PCC acting as a switch between internally and externally directed thoughts. This again supports research indicating the PCC is a major hub between networks^75–77^, particularly regarding task switching^83^. This may also help explain the inconsistent results previously mentioned regarding the role of the PCC at rest and during tasks, as if the role of the PCC is to shift between internally and externally directed thoughts, its coupling with each network will differ depending on the task at hand.

## Conclusions

We provide evidence for the lack of an effect of anodal tDCS over the left DLPFC in modulating mind-wandering propensity. Furthermore, we demonstrate a lack of an effect on brain activity and connectivity (both dynamic functional and effective) in three intrinsic brain networks thought to be associated with internal and externally directed thought processes (DMN, ECN, SN). This challenges previous research and highlights the need for replication studies.

## Acknowledgments

This work was supported by the Medical Research Council (MRC) via studentship to SC and partially by an MRC-NIRG grant (MR/P02596X/1) held by DF-E. We would also like to thank the Centre for Human Brain Health (CHBH), Birmingham, UK for providing in-kind access to MRI scanner time.

## Data Availability

The datasets generated during and/or analysed during the current study are available from the corresponding author on reasonable request.

